# Evaluation of cerebral and hepatic oxidative metabolism by administration of risperidone in a subacute model in rats (*Rattus norvegicus*)

**DOI:** 10.1101/2025.07.14.664632

**Authors:** Luis Salcedo-Valdez, Silvia Suárez-Cunza

**Affiliations:** Universidad Nacional Mayor de San Marcos, Facultad de Medicina San Fernando, Instituto de Investigación de Bioquímica y Nutrición (IIBN), Lima, Perú

**Author notes:** Corresponding author: Luis Salcedo-Valdez. Universidad Nacional Mayor de San Marcos, Facultad de Medicina San Fernando, Instituto de Investigación de Bioquímica y Nutrición (IIBN). Av. Miguel Grau 755, Lima 15001, Perú.

**Keywords:** Risperidone, oxidative stress, reactive oxygen species, antioxidants, lipid peroxidation

## Abstract

Risperidone is a second-generation antipsychotic widely prescribed for a variety of psychiatric disorders. Despite its widespread use, its subacute effects on oxidative metabolism in brain and liver tissues remain poorly understood. This study aimed to evaluate the impact of risperidone on antioxidant enzyme activity and lipid peroxidation in rats. A total of fifteen male Holtzman albino rats were randomly assigned to a Control group (n=5, no risperidone) and two treatment groups (n=5 per group) receiving 0.4 mg kg^-1^ day^-1^ and 4.0 mg kg^-1^ day^-1^ risperidone, administered via orogastric gavage for 20 consecutive days. After treatment, brain and liver tissues were collected. The activity of Superoxide Dismutase (SOD), Catalase (CAT), Glutathione Peroxidase (GPx), Glucose-6-Phosphate Dehydrogenase (G6PDH), and Glutathione S-Transferase (GST) was analyzed. Reduced Glutathione (GSH) levels and lipid peroxidation, measured as thiobarbituric acid reactive substances (TBARS), were quantified. Findings indicate that in brain tissue, both doses significantly increased CAT activity and decreased the SOD/CAT ratio, and that the high dose significantly reduced TBARS levels. In liver tissue, a significant increase in CAT activity was observed with the high dose. Furthermore, both doses significantly increased G6PDH activity and reduced TBARS levels. These results underscore the influence of risperidone on cerebral and hepatic oxidative metabolism during the subacute phase.

## Introduction

The prescription of antipsychotic medications has increased in response to the growing prevalence of mental health disorders. Among the most widely prescribed and commercially available antipsychotics is risperidone, a second-generation synthetic compound used in the long-term management of severe psychiatric conditions such as schizophrenia, as well as in the treatment of dementia, mood disorders, and autism (Turner 2020). Risperidone exerts its pharmacological effects primarily by antagonizing several neuronal receptors, including dopamine (D2) and serotonin (5-HT2), with higher affinity for the latter (Janssen et al. 1988, Leysen et al. 1994).

By blocking these receptors, risperidone disrupts intracellular dopamine and serotonin signaling, which can lead to altered monoamine metabolism and increased Monoamine Oxidase (MAO) activity, subsequently elevating hydrogen peroxide (H_2_O_2_) levels. In parallel, dopamine autoxidation may contribute to the generation of Reactive Oxygen Species (ROS), including superoxide anions (O_2_^•−^) and semiquinones (Muñoz et al. 2012, Halliwell & Gutteridge 2015). Moreover, Cytochrome P450 (CYP)-mediated metabolism of risperidone within neurons also produces ROS, further contributing to oxidative stress. In general, the accumulation of these chemical compounds could increase the amount of ROS, and, in turns, could promote cellular damage and alter the integrity of nervous tissues.

The brain is particularly vulnerable to oxidative damage due to its high lipid content, elevated metabolic rate, and disproportionately high oxygen consumption. However, it possesses a robust antioxidant defense system—including enzymes such as Superoxide Dismutases (Mn-SOD, CuZn- SOD), Glutathione Peroxidase (GPx), Catalase (CAT), Glutathione S-Transferase (GST), and metabolites like Reduced Glutathione (GSH)—however, these defenses may be insufficient under sustained oxidative stress (Halliwell & Gutteridge 2015).

The liver, a critical organ in drug metabolism, plays a central role in the detoxification of risperidone. Hepatic metabolism involves extensive antioxidant and enzymatic activity. More than 1000 drugs have been implicated in hepatic side effects, with neuropsychiatric drugs accounting for approximately 16% of cases (Dumortier et al. 2002). Risperidone has been associated with hepatotoxicity during long-term use, including hepatocellular damage, altered liver enzyme profiles, and cholestatic hepatitis (Krebs et al. 2001, Kumra et al. 1997, Lopez-Torres et al. 2014, Mouradian-Stamatiadis et al. 2002). Oxidative stress, driven by increased ROS production, is believed to underlie much of this hepatic damage (Eftekhari et al. 2016).

Despite its widespread clinical use, the subacute effects of risperidone on oxidative metabolism in both brain and liver tissues remain incompletely characterized. Therefore, in this study, we evaluated the activity of key antioxidant enzymes in control rats and those that received risperidone at two different doses for 20 days—an exposure period equivalent to approximately 16 months in humans (Pillai et al. 2007).

## Materials and Methods

### Animal Protocol

Fifteen male Holtzman albino rats (initial body weight: 245 ± 13 g) were obtained from La Molina National Agrarian University. Animals were housed individually in metal cages at the bioterium of the Faculty of Medicine, San Marcos University. Standard housing conditions were maintained: temperature at 24 ± 2 °C, relative humidity at 60 ± 5 %, and a 12-h light/dark cycle. Rats had *ad libitum* access to food and water and were acclimatized for seven days prior to experimentation.

### Chemicals

Risperidone was obtained as 2 mg Zeclonex™ tablets. For its preparation, the tablets were crushed and dissolved in bidistilled water. Reagents purchased from Sigma-Aldrich: 4,6- dihydroxypyrimidine-2-thiol (thiobarbituric acid, TBA), potassium chloride, sodium chloride, disodium phosphate, monopotassium phosphate, benzene-1,2,3-triol (pyrogallol), 2-amino-2- (hydroxymethyl)propane-1,3-diol (Tris base), 2-amino-2-(hydroxymethyl)propane-1,3-diol hydrochloride (Tris-HCl), diethylenetriaminepentaacetic acid (DTPA), magnesium chloride, β- nicotinamide adenine dinucleotide phosphate (oxidized; NADP^+^), glucose-6-phosphate, ethylenediaminetetraacetic acid (EDTA), reduced glutathione (GSH), β-nicotinamide adenine dinucleotide phosphate (reduced; NADPH), glutathione reductase, 1-chloro-2,4-dinitrobenzene (CDNB), 5,5′-dithiobis-(2-nitrobenzoic acid) (DTNB), and bovine serum albumin. Additional reagents included trichloroacetic acid (TCA), hydrogen peroxide (H_2_O_2_), absolute ethanol, copper (II) sulfate, and sodium hydroxide (Merck); hydrochloric acid (Fisher Scientific); and sodium azide (NaN_3_) (Fluka).

### Experimental Design

Fifteen male Holtzman rats were randomly divided into three groups (*n* = 5) using the OpenEpi program. The groups were: Control (received 2 mL of bidistilled water), RISP1 (0.4 mg kg^-1^ day^-1^ risperidone), and RISP2 (4.0 mg kg^-1^ day^-1^ risperidone). Doses were selected based on previous studies using risperidone. All administrations (2 mL) were performed via orogastric gavage once daily between 16:00 and 17:00 h for 20 consecutive days. At the end of treatment, the animals were euthanized and brain and liver tissues were immediately collected and stored at 4 °C.

### Tissue Homogenate Preparation

Excised organs were perfused with cold 0.154 mol L^-1^ KCl to remove blood, blotted, and weighed. Tissue homogenates (10 % w/v) were prepared in phosphate-buffered saline (PBS) using a Potter-Elvehjem homogenizer with a Teflon pestle. Homogenates were centrifuged at 7267 m s^-2^ (741 × *g*, where *g* is the standard acceleration due to gravity) for 5 min at 4 °C. (MPW Med instruments, MPW380R). Supernatants were collected and stored on ice. All biochemical analyses were completed within 48 h of tissue preparation and carried out in an ice bath using a spectrophotometer (Thermo Fisher Scientific G10S UV-Vis).

### Superoxide Dismutase (SOD) Activity

SOD activity was measured using the pyrogallol autoxidation inhibition method (Marklund & Marklund 1974). A control reaction for spontaneous autoxidation was established with 950 μL of 50 mmol L^-1^ Tris-HCl buffer (pH 8.2) containing 1 mmol L^-1^ DTPA. After a 1-min incubation at 37 °C, 50 μL of 2 mmol L^-1^ pyrogallol (prepared in 10 mmol L^-1^ HCl) was added. For enzyme activity measurements, 50 μL of tissue homogenate replaced an equivalent volume of buffer. The rate of absorbance change was recorded over 3 min at 420 nm. One unit (U) of SOD activity was defined as the amount of enzyme required to produce a 50 % inhibition in the rate of pyrogallol autoxidation, corresponding to an absorbance change of 0.02 per minute (±10 %).

### Catalase (CAT) Activity

CAT activity was determined following the method of Aebi (1984). In a 1 mL quartz cuvette, 0.6 mL of tissue homogenate was combined with 0.3 mL of 30 mmol L^-1^ H_2_O_2_ prepared in 50 mmol L^-1^ phosphate buffer pH 7.0. The decomposition of H_2_O_2_ was monitored by measuring the decrease in absorbance at 240 nm over 2 min using a spectrophotometer. Enzyme activity was calculated using a molar extinction coefficient of 43.6 L mol^−1^ cm^−1^.

### Glutathione Peroxidase (GPx) Activity

GPx activity was measured using the protocol of Paglia and Valentine (1967). A reaction buffer was prepared consisting of 0.1 mol L^-1^ phosphate buffer (pH 7.0), 1 mmol L^-1^ EDTA, 4 mmol L^-1^ NaN_3_, 6.1 mg mL^-1^ GSH, 10 mmol L^-1^ NADPH, and 1000 U mL^-1^ glutathione reductase. In a 1 mL cuvette, 980 µL of the substrate buffer and 10 µL of homogenate were mixed and incubated at 25 °C for 5 min. The reaction was initiated by adding 10 µL of 8.8 mmol L^-1^ H_2_O_2_, and the decrease in absorbance was monitored at 340 nm for 3 min. GPx activity was calculated using the molar extinction coefficient of 6220 L mol^−1^ cm^−1^.

### Glucose-6-Phosphate Dehydrogenase (G6PDH) Activity

G6PDH activity was assessed according to Kjeldsberg et al. (1989). In a 1 mL spectrophotometric cuvette, the following were added: 200 µL of 50 mmol L^-1^ Tris-HCl buffer (pH 8.8), 160 µL of 0.1 mol L^-1^ MgCl_2_, 40 µL of 6.0 mmol L^-1^ NADP^+^, 40 µL of tissue homogenate, and 720 µL of double-distilled water. After incubating the mixture at 30 °C for 2 min, the reaction was initiated by adding 40 µL of 36 mmol L^-1^ glucose-6-phosphate. The increase in absorbance was recorded at 340 nm for 2 min. Enzyme activity was expressed using the molar extinction coefficient of 6220 L mol^−1^ cm^−1^.

### Glutathione S-Transferase (GST) Activity

GST activity was determined using the method described by Habig et al. (1974). The reaction mixture contained 100 mmol L^-1^ CDNB in ethanol, 100 mmol L^-1^ GSH, and 0.1 mol L^-1^ phosphate buffer (pH 6.5). In a 1 mL cuvette, 980 µL of this reagent mix was incubated at 25 °C for 3 min. Subsequently, 20 µL of homogenate was added, and the increase in absorbance was measured at 340 nm for 3 min. Enzymatic activity was calculated using a molar extinction coefficient of 9.6 L mmol^−1^ cm^−1^.

### Reduced Glutathione (GSH) Quantification

GSH concentration was quantified following the method of Boyne and Ellman (1972). Briefly, 950 µL of tissue homogenate was mixed with 50 µL of 100 % (w/v) TCA and centrifuged at 74371 m s^-2^ (7585 × *g*) for 15 min. Then, 200 µL of the supernatant was added to 1 mL of 0.5 mol L^-1^ potassium phosphate buffer (pH 6.8) and incubated at room temperature (25 °C) for 25 min. Subsequently, 200 µL of DTNB (1.5 mg mL^−1^ in the same potassium phosphate buffer) was added, and the mixture was incubated for an additional 5 min. Absorbance was measured at 412 nm. A 4 mmol L^-1^ GSH standard, diluted 1:10 in 5 % (w/v) TCA, was used for calibration.

### Lipid Peroxidation Assay

Lipid peroxidation levels were determined using the method described by Buege and Aust (1978). Briefly, 0.3 mL of brain or liver homogenate was mixed with 0.6 mL of 10 % or 20 % (w/v) TCA, respectively, and incubated in a boiling water bath for 15 min. After cooling under running tap water, 0.67 % TBA (w/v) in 0.25 mol L^-1^ HCl was added. Samples were reheated for 30 min in a boiling water bath, cooled with cold water, and centrifuged at 56948 m s^-2^ (5807 × *g*) for 10 min at room temperature (25 °C). The absorbance of the supernatant was measured at 535 nm.

### Total Protein Quantification

Total protein content in tissue homogenates was determined using the Biuret method (Gornall et al. 1949). The reaction mixture consisted of 150 µL of double-distilled water, 150 µL of 40 mmol L^-1^ copper (II) sulfate, 100 µL of tissue homogenate, and 1 mL of 2.5 mmol L^-1^ sodium hydroxide. A 2 % (w/v) bovine serum albumin solution was used as a standard. After 5 min of incubation at room temperature (25 °C), absorbance was read at 540 nm.

### Statistical Analysis

All data were expressed as mean ± standard deviation (SD). Statistical comparisons among groups were conducted using one-way analysis of variance (ANOVA) followed by Tukey’s *post hoc* test for multiple comparisons. Analyses were performed using R (version 4.3.2) and GraphPad Prism (version 8). Differences were considered statistically significant at *p* < 0.05.

### Ethical Aspects

Experimental procedures were carried out in accordance with the Guide for the Care and Use of Laboratory Animals (National Research Council 2011) and the ARRIVE Essential Guidelines (Percie du Sert et al. 2020). The design and execution of this research rigorously adhered to the 3Rs principles (Replacement, Reduction, and Refinement) for animal welfare. All necessary measures were implemented to ensure animal well-being and minimize suffering throughout the experiment, as stated in Law No. 30407, the Animal Protection and Welfare Law.

## Results

In brain tissue homogenates, a significant increase in CAT activity was observed (Table 1). On the other hand, the activities of SOD, GPx, and the SOD/GPx ratio showed no significant differences. The SOD/CAT ratio, however, did present a significant decrease (Table 1). G6PDH and GST activities, along with GSH levels, remained unchanged across all groups (Table 2). Regarding lipid peroxidation (TBARS), a significant reduction was observed in the high-dose risperidone group (RISP2, 4.0 mg kg^-1^ day^-1^) compared to the Control and RISP1 groups (Figure 1).

**Table 1.** Effect of subacute risperidone treatment on the specific activity of the antioxidant enzymes SOD, CAT and GPx in rat brain tissue.

| Parameters | Groups |  |  |
| --- | --- | --- | --- |
|  | Control | RISP1 | RISP2 |
| Dose (mg kg<sup>-1</sup> day<sup>-1</sup>) | - | 0.4 | 4.0 |
| SOD (U mg<sup>-1</sup> protein) | 5.79 ± 0.2 | 6.07 ± 0.7 | 5.60 ± 0.7 |
| CAT (μmol H<sub>2</sub>O<sub>2</sub> min<sup>-1</sup> mg<sup>-1</sup> protein) | 4.42 ± 0.1<sup>a</sup> | 7.27 ± 0.6<sup>b</sup> | 6.50 ± 1.1<sup>b</sup> |
| GPx (nmol NADPH min<sup>-1</sup> mg<sup>-1</sup> protein) | 67.7 ± 7.6 | 73.5 ± 6.7 | 68.1 ± 3.5 |
| SOD/CAT | 1.31 ± 0.06<sup>a</sup> | 0.84 ± 0.11<sup>b</sup> | 0.88 ± 0.19<sup>b</sup> |
| SOD/GPx | 0.09 ± 0.01 | 0.08 ± 0.01 | 0.08 ± 0.01 |

RISP: risperidone, SOD: superoxide dismutase, U: unit of enzymatic activity, CAT: catalase, GPx: glutathione peroxidase. Risperidone (0.4 and 4.0 mg kg<sup>-1</sup> day<sup>-1</sup> treatments) was prepared in bidistilled water and administered in a volume of 2 mL orogastrically to male Holtzman rats (*n* = 5 per group) for 20 days. The Control group received 2 mL of bidistilled water. Results are presented as mean ± SD. Different lowercase letters within a row indicate statistically significant differences (*p* < 0.05) according to Tukey's *post hoc* test.

**Table 2.** Effect of subacute risperidone treatment on the specific activity of the antioxidant enzymes G6PDH and GST and the concentration of GSH in rat brain tissue.

| Parameters | Groups |  |  |
| --- | --- | --- | --- |
|  | Control | RISP1 | RISP2 |
| Dose (mg kg<sup>-1</sup> day<sup>-1</sup>) | - | 0.4 | 4.0 |
| G6PDH (nmol NADPH min<sup>-1</sup> mg<sup>-1</sup> protein) | 31.9 ± 2.7 | 30.0 ± 3.2 | 28.4 ± 2.9 |
| GST (mmol CDNB conjugated min<sup>-1</sup> mg<sup>-1</sup> protein) | 53.9 ± 3.2 | 56.7 ± 6.1 | 54.1 ± 6.5 |
| GSH (nmol GSH mL<sup>-1</sup> homogenate) | 90.5 ± 9.8 | 85.5 ± 12.2 | 88.6 ± 13.1 |

RISP: risperidone, G6PDH: glucose-6-phosphate dehydrogenase, GST: glutathione S-transferase, GSH: reduced glutathione. Risperidone (0.4 and 4.0 mg kg<sup>-1</sup> day<sup>-1</sup> treatments) was prepared in bidistilled water and administered in a volume of 2 mL orogastrically to male Holtzman rats ( $n = 5$ per group) for 20 days. The Control group received 2 mL of bidistilled water. Results are presented as mean ± SD. No significant differences were found between groups ( $p > 0.05$ ) according to one-way ANOVA.

**Figure 1.**
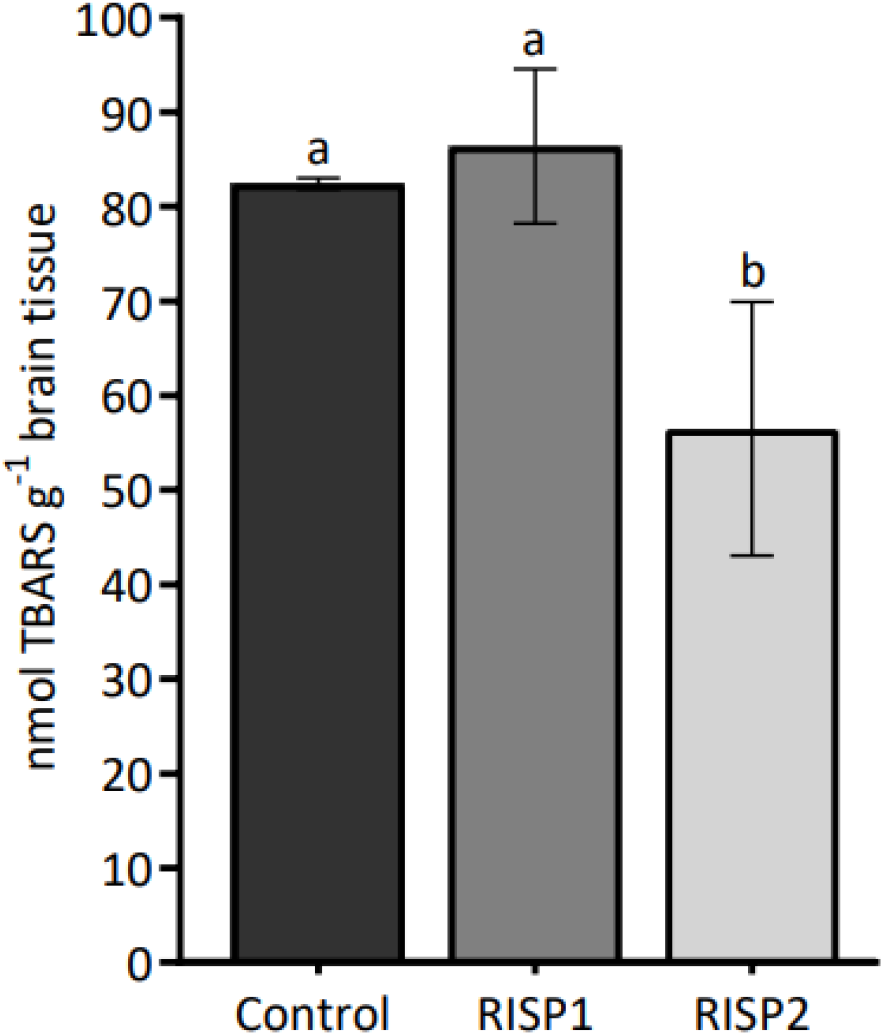
Levels of lipid peroxidation in brain tissue of male Holtzman rats treated with risperidone. TBARS: thiobarbituric acid reactive substances. Risperidone doses and bidistilled water were prepared in a volume of 2 mL of bidistilled water and administered by orogastric gavage. Control: received 2 mL of bidistilled water (*n* = 5), RISP1: 0.4 mg kg^−1^ day^−1^ risperidone in 2 mL (*n* = 5), RISP2: 4.0 mg kg^-1^ day^-1^ risperidone in 2 mL (*n* = 5). Rats were treated subacutely (20 days). TBARS values are presented as mean ± SD. Different lowercase letters between bars indicate significant differences (*p* < 0.05) according to Tukey’s *post hoc* test.

In liver tissue homogenates, CAT activity significantly increased in the RISP2 group (4.0 mg kg^-1^ day^-1^) compared to the Control group (Table 3). Conversely, SOD and GPx activities, along with the SOD/CAT and SOD/GPx ratios, showed no significant changes (Table 3). G6PDH activity significantly increased with both risperidone doses (0.4 and 4.0 mg kg^-1^ day^-1^), while GST activity and GSH levels remained unchanged (Table 4). TBARS levels showed a significant decrease in both risperidone-treated groups (RISP1: 0.4 mg kg^-1^ day^-1^ and RISP2: 4.0 mg kg^-1^ day^-1^) compared to the Control (Figure 2).

**Table 3.** Effect of subacute risperidone treatment on the specific activity of the antioxidant enzymes SOD, CAT and GPx in rat liver tissue.

| Parameters | Groups |  |  |
| --- | --- | --- | --- |
|  | Control | RISP1 | RISP2 |
| Dose (mg kg<sup>-1</sup> day<sup>-1</sup>) | - | 0.4 | 4.0 |
| SOD (U mg<sup>-1</sup> protein) | 12.2 ± 1.7 | 13.3 ± 0.9 | 13.6 ± 0.9 |
| CAT (μmol H<sub>2</sub>O<sub>2</sub> min<sup>-1</sup> mg<sup>-1</sup> protein) | 292 ± 30<sup>a</sup> | 315 ± 25<sup>ab</sup> | 338 ± 23<sup>b</sup> |
| GPx (nmol NADPH min<sup>-1</sup> mg<sup>-1</sup> protein) | 281 ± 60 | 353 ± 50 | 302 ± 55 |
| SOD/CAT | 0.04 ± 0.01 | 0.04 ± 0.01 | 0.04 ± 0.01 |
| SOD/GPx | 0.04 ± 0.004 | 0.04 ± 0.01 | 0.05 ± 0.01 |

RISP: risperidone, SOD: superoxide dismutase, U: unit of enzymatic activity, CAT: catalase, GPx: glutathione peroxidase. Risperidone (0.4 and 4.0 mg kg<sup>-1</sup> day<sup>-1</sup> treatments) was prepared in bidistilled water and administered in a volume of 2 mL orogastrically to male Holtzman rats (*n* = 5 per group) for 20 days. The Control group received 2 mL of bidistilled water. Results are presented as mean ± SD. Different lowercase letters within a row indicate statistically significant differences (*p* < 0.05) according to Tukey's *post hoc* test.

**Table 4.** Effect of subacute risperidone treatment on the specific activity of the antioxidant enzymes G6PDH, GST and the concentration of GSH in rat liver tissue.

| Parameters | Groups |  |  |
| --- | --- | --- | --- |
|  | Control | RISP1 | RISP2 |
| Dose (mg kg<sup>-1</sup> day<sup>-1</sup>) | - | 0.4 | 4.0 |
| G6PDH (nmol NADPH min<sup>-1</sup> mg<sup>-1</sup> protein) | 4.1 ± 0.7<sup>a</sup> | 7.4 ± 1.8<sup>b</sup> | 9.6 ± 2.0<sup>b</sup> |
| GST (mmol CDNB conjugated min<sup>-1</sup> mg<sup>-1</sup> protein) | 320 ± 7.7 | 333 ± 37 | 321 ± 33 |
| GSH (nmol GSH mL<sup>-1</sup> homogenate) | 548 ± 111 | 516 ± 59 | 548 ± 24 |

RISP: risperidone, G6PDH: glucose-6-phosphate dehydrogenase, GST: glutathione S-transferase, GSH: reduced glutathione. Risperidone (0.4 and 4.0 mg kg<sup>-1</sup> day<sup>-1</sup> treatments) was prepared in bidistilled water and administered in a volume of 2 mL orogastrically to male Holtzman rats (*n* = 5 per group) for 20 days. The Control group received 2 mL of bidistilled water. Results are presented as mean ± SD. Different lowercase letters within a row indicate statistically significant differences (*p* < 0.05) according to Tukey's *post hoc* test.

**Figure 2.**
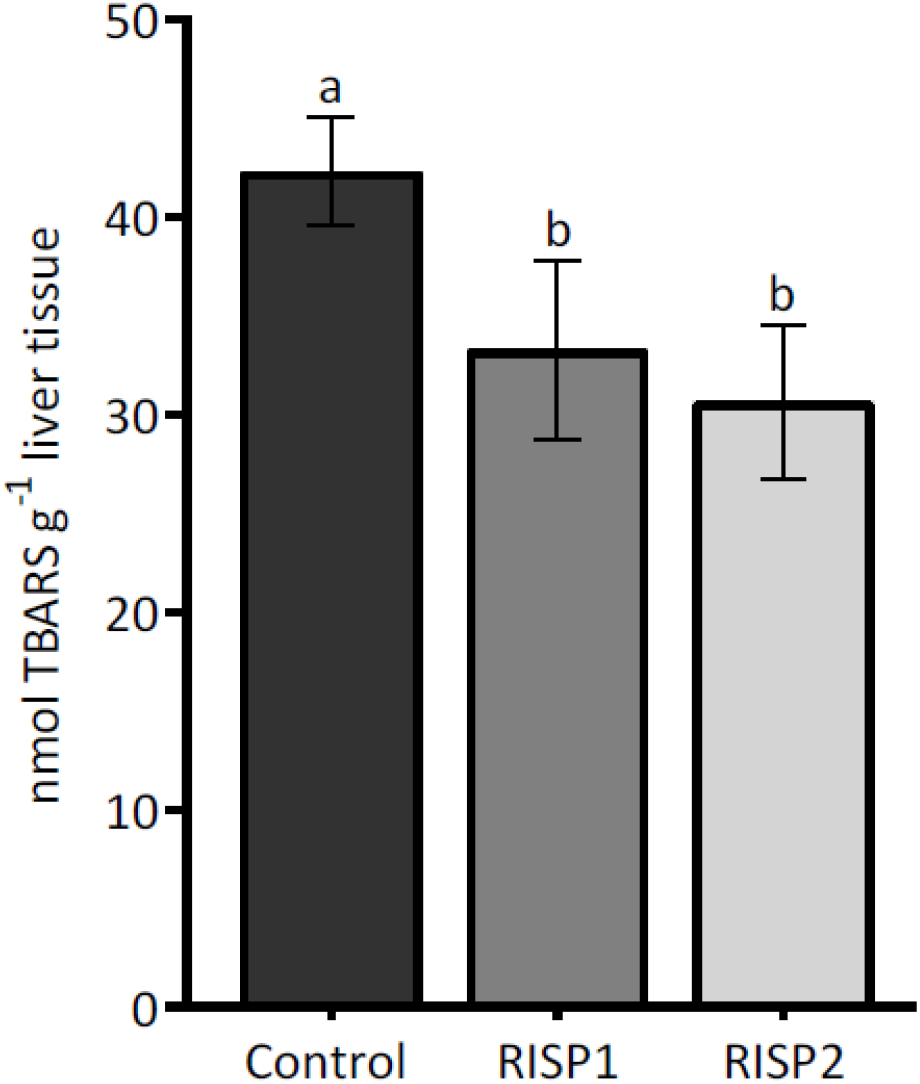
Levels of lipid peroxidation in liver tissue of male Holtzman rats treated with risperidone. TBARS: thiobarbituric acid reactive substances. Risperidone doses and bidistilled water were prepared in a volume of 2 mL of bidistilled water and administered by orogastric gavage. Control: received 2 mL of bidistilled water (*n* = 5), RISP1: 0.4 mg kg^-1^ day^-1^ risperidone in 2 mL (*n* = 5), RISP2: 4.0 mg kg^-1^ day^-1^ risperidone in 2 mL (*n* = 5). Rats were treated subacutely (20 days). TBARS values are presented as mean ± SD. Different lowercase letters between bars indicate significant differences (*p* < 0.05) according to Tukey’s *post hoc* test.

Lastly, total protein concentrations in both brain and liver homogenates showed no statistically significant variation across experimental groups.

## Discussion

The findings demonstrate that subacute risperidone administration over 20 days affects redox homeostasis in rat brain and liver tissues, albeit in tissue-specific and enzyme-selective ways. In brain homogenates, CAT activity significantly increased at both risperidone doses, 0.4 mg kg^-1^ day^-1^ and 4.0 mg kg^-1^ day^-1^ (Table 1). CAT is responsible for degrading H_2_O_2_, which is produced as a byproduct of several cellular processes. One major source of H_2_O_2_ is the dismutation of O_2_^•−^ catalyzed by SOD. Additionally, H_2_O_2_ can be generated through the cytochrome P450 2D6 (CYP2D6)-mediated metabolism of risperidone to 9-hydroxyrisperidone. Furthermore, risperidone’s antagonism of dopamine and serotonin receptors leads to increased synaptic concentrations of these neurotransmitters, which may undergo oxidative deamination by MAO, producing H_2_O_2_ as a metabolic byproduct. The convergence of these pathways likely contributes to the observed upregulation of CAT activity as a compensatory mechanism to metabolize the excess H_2_O_2_ formed.

Thus, these diverse pathways of H_2_O_2_ production might also explain the non-significant results for GPx, another key H_2_O_2_-detoxifying enzyme (Table 1). GPx possesses a lower Km for H_2_O_2_ than CAT, indicating it is typically more active at lower substrate concentrations. The lack of significant GPx activation in this study may indicate that H_2_O_2_ levels exceeded the range where GPx is most effective, potentially saturating or even inhibiting the enzyme. Indeed, high concentrations of H_2_O_2_ have been reported to inhibit GPx activity during prolonged oxidative stress (Pigeolet et al. 1990, Cho et al. 2010). However, 20-day treatment period may have been insufficient to reveal this effect.

In contrast, Stojković et al. (2012) reported a decrease in GPx activity in thalamic tissue homogenate after 65 days of treatment with 0.84 mg kg^-1^ day^-1^ risperidone. However, Parikh et al. (2003) and Pillai et al. (2007) found no significant changes in GPx activity in brain homogenates after treatments of 45 and 90 days, and 90 and 180 days of treatment, respectively, both using a dose of 2.5 mg kg^-1^ day^-1^. Surprisingly, in a short 14-day treatment with 1.2 mg kg^-1^ risperidone in tissue homogenates from the prefrontal cortex, hippocampus, caudate nucleus, putamen, and amygdala, risperidone did not alter GPx activity (Casquero-Veiga et al. 2019). These mixed outcomes regarding GPx activity following risperidone administration, along with our own findings, underscore that enzyme response likely depends on crucial factors such as treatment duration, brain region, and dosage.

The absence of significant changes in SOD activity may be attributed to the intrinsic antioxidant properties of risperidone itself (Table 1). This suggests that O_2_^•−^ may be neutralized not only through enzymatic dismutation by SOD but also via radical dissipation through the antioxidant capacity of risperidone. The chemical structure of risperidone contains functional groups capable of stabilizing unpaired electrons, potentially enabling it to capture and neutralize O_2_^•−^ independently of enzymatic mechanisms (Brinholi et al. 2016).

Similar results to those mentioned in catalase studies, may apply to SOD activity under extended risperidone exposure. In that long-term study of Pillai et al. (2007), it was observed that at 180 days the activity of this enzyme decreased significantly in rats receiving treatment; however, SOD activity is preserved up to 90 days of treatment (Pillai et al. 2007). Risperidone may still be preserving its antioxidant activity.

Our results further support the notion of a redox imbalance in brain tissue, as evidenced by a significant decrease in the SOD/CAT ratio in both risperidone-treated groups compared to the Control (Table 1). This shift toward CAT activity suggests that the source of H_2_O_2_ extends beyond SOD-mediated superoxide dismutation. As previously discussed, H_2_O_2_ is also generated through microsomal risperidone metabolism via CYP2D6 and through MAO activity.

In terms of non-enzymatic antioxidant defense, neither significant changes were observed in brain GSH levels nor in the activity of G6PDH, an enzyme related to levels of this tripeptide. Neither was a significant change observed in the activity of GST, whose cosubstrate is GSH (Table 2). Thus, at the brain level, GSH-related oxidative metabolism is not affected by risperidone administration.

Regarding lipid peroxidation, a significant reduction was observed in TBARS levels in the brain following risperidone treatment at 4.0 mg kg^-1^ day^-1^ (Figure 1). Specifically, the results show a decrease of 31 % in the 4.0 mg kg^-1^ day^-1^ group compared to the Control. This reduction, alongside increased CAT activity, points toward a protective antioxidant effect of risperidone under subacute exposure conditions. The upregulation of CAT may have accelerated H_2_O_2_ decomposition, thereby mitigating the formation of hydroxyl radicals that initiate lipid peroxidation.

Our findings align with those of Stojković et al. (2012), who reported a significant decrease in malondialdehyde (MDA) levels in the cerebral cortex after 65 days of risperidone administration (0.84 mg kg^-1^ day^-1^). This finding supports the idea that under certain conditions this drug may have protective effects against lipid peroxidation in the cerebral cortex, although there were no changes in other brain regions. In other studies, using a 2.5 mg kg^-1^ day^-1^ dose, no significant changes were found after 45 days (Parikh et al. 2003) and 90 days of treatment (Parikh et al. 2003, Pillai et al. 2007). Only a 180-day treatment with 2.5 mg kg^-1^ day^-1^ dose triggered an increase in lipoperoxidation (Pillai et al. 2007), a peroxidative process that can be initiated by different types of ROS.

On the other hand, the liver exhibits a distinct oxidative and antioxidant response profile, largely attributable to its central role in systemic detoxification. Under the conditions evaluated in this study, while SOD activity did not show significant alterations in liver homogenates following subacute risperidone administration, CAT activity significantly increased in the RISP2 group (4.0 mg kg^-1^ day^-1^) compared to the Control group (Table 3). It is possible that the O_2_^•−^ generated during hepatic risperidone metabolism—primarily via the CYP2D6 enzymatic pathway—was either effectively neutralized by the liver’s robust antioxidant defenses, rapidly reacted with organic substrates, or was simply produced in quantities too low to provoke measurable enzymatic responses that would affect SOD. Despite SOD activity remaining unaffected, the observed increase in CAT activity at the higher dose suggests a compensatory response to a heightened H_2_O_2_ burden, which could originate from risperidone’s hepatic metabolism (including CYP2D6 activity and other pathways) or from the basal action of SOD.

Nonetheless, a significant increase in G6PDH activity was observed at both the lower (0.4 mg kg^- 1^ day^-1^) and higher (4.0 mg kg^-1^ day^-1^) doses of risperidone compared to the Control group (Table 4). This increase suggests an increased demand for reducing equivalents in the form of NADPH + H^+^. NADPH + H^+^ is essential for maintaining GSH in its reduced form and thereby sustaining the activity of GSH-dependent enzymes such as GPx and GST, both of which, importantly, showed no significant changes in our study (Table 4). The elevated G6PDH activity, thus, highlights the liver’s potential for risperidone detoxification by ensuring adequate supply of NADPH + H^+^. Additionally, this reducing equivalent is also required for CYP2D6 activity, which is involved in risperidone metabolism. The significant increase in G6PDH activity at both doses supports the notion of an enhanced need for NADPH + H^+^. to fuel these detoxification pathways.

Our findings contrast with those of Eftekhari et al. (2016), who reported a significant reduction in hepatic GSH levels following an acute, single high-dose (6 mg kg^-1^) administration of risperidone, assessed just 5 h post-treatment. Our results showed no significant changes in GSH levels after 20-day treatment with any doses we applied. This discrepancy highlights the importance of dose and exposure duration in modulating hepatic antioxidant responses. Although risperidone may exert an initial acute effect on GSH, the hepatic system may develop an adaptive or compensatory mechanism over the course of a 20-day subacute exposure, thereby managing to maintain its levels of this crucial antioxidant and its detoxification capacity. These mechanisms involved in oxidative metabolism resulted in a significant decrease in lipid peroxidation, as evidenced by reduced TBARS levels in both risperidone-treated groups compared to the Control group (Figure 2). Specifically, our results show a decrease in lipid peroxidation of approximately 21 % in the 0.4 mg kg^-1^ day^-1^ group and 27 % in the 4.0 mg kg^-1^ day^-1^ group compared to Control. These findings suggest that the antioxidant defense mechanisms in the liver were sufficient or were activated to actively mitigate lipid damage under the conditions studied. However, this contrasts with evidence from Eftekhari et al. (2016), who observed a significant increase in MDA following a single 6 mg kg^-1^ dose over a short time frame. Thus, risperidone’s prooxidant effects may be appreciated at concentrations or exposure times different from those investigated in our study. Then, considering the antioxidant activities of enzymes independent of GSH—SOD and CAT—and the enzymes associated with GSH levels, it can be observed that the effects of risperidone on antioxidant defense metabolisms differ between the liver and the brain.

Overall, 20-day risperidone treatments at both 0.4 mg kg^-1^ day^-1^ and 4.0 mg kg^-1^ day^-1^ significantly increased cerebral CAT enzymatic activity in male Holtzman rats. Moreover, the higher dose (4.0 mg kg^-1^ day^-1^) also led to a significant decrease in cerebral TBARS, suggesting a rapid protective response to increased H_2_O_2_ generation linked to risperidone metabolism. In the liver tissue, at the higher dose (4.0 mg kg^-1^ day^-1^), we observed a significant increase in hepatic CAT activity, which plays a crucial role in detoxifying H_2_O_2_. Furthermore, both the lower (0.4 mg kg^-1^ day^-1^) and higher (4.0 mg kg^-1^ day^-1^) doses of risperidone promoted a significant increase in G6PDH enzymatic activity. This increase is crucial as it ensures an enhanced supply of NADPH + H^+^, which is vital for both CYP2D6-mediated risperidone metabolism and the regeneration of reduced GSH, thus supporting GSH-dependent enzyme systems. Collectively, these changes in enzymatic activities, along with the significant decrease in TBARS levels at both risperidone doses, suggest a robust and effective hepatic antioxidant defense mechanism actively mitigating oxidative damage.

Thus, the findings of this study provide valuable information on both tissues, liver and brain, during subacute treatment, a time period that is not usually considered significant by the effects of a drug typically used in long-term treatments. Finally, further studies are needed to explore additional aspects of the detoxification process, particularly those involving Phase II and III metabolism and the transcriptional regulation of antioxidant defenses. Such investigations may provide a more comprehensive view of risperidone’s metabolic footprint from the onset of therapy onward.

## Rol de los autores / Authors Roles

LSV: conceptualization, formal analysis, investigation, writing – original draft, writing – review and editing, visualization.

SSC: conceptualization, methodology, investigation, resources, writing – review and editing, visualization, project administration, supervision, funding acquisition.

## Conflicto de intereses / Competing interests

The authors declare no conflict of interest.

## Fuentes de financiamiento / Funding

Funding for this research was provided by PCONFIGI 150104161, with the approval of the VRIP of the National University of San Marcos (Universidad Nacional Mayor de San Marcos).

## Agradecimientos / Acknowledgments

We thank Medical Technologist Oswaldo Sanabria Salazar for his support in determining the enzymatic activity of catalase and the lipid peroxidation assay.

## Aspectos éticos / legales; Ethics / legals

The authors declare that they have not violated or omitted any ethical or legal norms in conducting this research and producing this work.

